# Integrated biophysical characterization of fibrillar collagen-based hydrogels

**DOI:** 10.1101/868059

**Authors:** Alex Avendano, Jonathan C. Chang, Marcos G. Cortes-Medina, Aaron J. Seibel, Bitania R. Admasu, Cassandra M. Boutelle, Andrew R. Bushman, Ayush Arpit Garg, Cameron M. DeShetler, Sara L. Cole, Jonathan W. Song

## Abstract

This paper describes an experimental characterization scheme of the biophysical properties of reconstituted hydrogel matrices based on indentation testing, quantification of transport via microfluidics, and confocal reflectance microscopy analysis. While methods for characterizing hydrogels exist and are widely used, they often do not measure diffusive and convective transport concurrently, determine the relationship between microstructure and transport efficiency, and decouple matrix mechanics and transport properties. Our integrated approach enabled independent and quantitative measurements of the structural, mechanical, and transport properties of hydrogels in a single study. We used fibrillar type I collagen as the base matrix and investigated the effects of two different matrix modifications: 1) cross-linking with human recombinant tissue transglutaminase II (hrTGII) and 2) supplementation with the non-fibrillar matrix constituent hyaluronic acid (HA). hrTGII modified matrix structure and transport but not mechanical parameters. Furthermore, changes in matrix structure due to hrTGII were seen to be dependent on the concentration of collagen. In contrast, supplementation of HA at different collagen concentrations altered matrix microstructure and mechanical indentation behavior but not transport parameters. These experimental observations reveal the important relationship between ECM composition and biophysical properties. The integrated techniques are versatile, robust, and accessible; and as matrix-cell interactions are instrumental for many biological processes, the methods and findings described here should be broadly applicable for characterizing hydrogel materials used for 3-D tissue engineered culture models.

## 1. Introduction

The tissue space between living cells, commonly known as the interstitium, is comprised primarily of structural extracellular matrix (ECM) proteins and aqueous fluid. This environment is complex, information-rich, and provides cells with instructive cues to dictate emergent behavior^1^. ECM composition has a profound influence on the biophysical properties of the interstitium^2–4^ These properties include structural, mechanical, and transport, which in turn can directly or indirectly affect cell behavior and tissue-level response^5–7^. For instance, the predominant ECM protein *in vivo* is type I collagen, which forms a fibrillar network. The microstructural properties of the collagen network, such as mean pore size, fiber alignment, and fiber radius are known to influence cell adhesion, proliferation, migration, and differentiation^8–10^. This collagen scaffold can be further modified by cross-linking agents such as tissue transglutaminase 2 (TGII)^3, 11–12^. In addition to collagen, glycosaminoglycans (GAGs) are an important structural ECM constituent that helps provide hydrated gel-like properties of the interstitium^13^. The collagen and GAG ECM compositions can vary greatly by tissue-type and be altered dramatically during diseased states such as cancer and fibrosis^14^. These matrix alterations impact the mechanical compliance at both the bulk and local tissue levels and their subsequent effects on fundamental cell behaviors such as morphogenesis, proliferation, motility, differentiation, and response to therapeutic agents^15–16^. Mass transport properties of the interstitium also affect cell behavior by modulating shear stresses, biomolecular gradients, nutrient availability, waste clearance, and drug delivery^17–22^.

While the biophysical properties of tissue can be measured *in vivo*, studying these properties in this setting is challenging due to limited experimental control over ECM composition. With regards to transport properties, it is very difficult to specify concentration and pressure gradients *in vivo* and identify the independent contributions of diffusion and convection. In response, 3-D hydrogel-based culture models comprised of either synthetic or natural biomaterials have been widely adopted to provide physiological-like settings *in vitro*. Among the materials used to mimic the interstitial ECM, reconstituted type I collagen and the non-sulfated GAG hyaluronic acid (HA) have been used extensively due to their abundance in the interstitium, commercial availability, and versatility^23–24^. Characterization of collagen hydrogel systems in terms of structure, mechanics, and transport has been accomplished in the past^25^. Yet, these previous studies have not provided integrated analysis of these biophysical properties of collagen-based matrices in a single study, which among other challenges makes it difficult to simultaneously consider diffusive and convective transport, establish the relationship between microstructural parameters and transport efficiency, and decouple matrix mechanics and transport properties.

Detailed and systematic characterization of the biophysical properties of reconstituted acellular hydrogels is necessary in order to define the initial conditions for the dynamic and two-way reciprocal interactions between matrix and cells. These findings can benefit others in studying ECM reorganization in health and disease along with its subsequent impact on cell function. Therefore, the goal of this study is to present an integrated biophysical characterization scheme using mechanical indentation testing, confocal reflectance imaging and analysis, microfluidics, and fluorescence recovery after photobleaching (FRAP) to profile structural, mechanical, and transport properties of acellular collagen-based hydrogels. This methodology has the advantages of: 1) simultaneously integrating structure, mechanical, and transport profiling in a single study 2) independently investigating convective and diffusive transport, and 3) decoupling matrix mechanics and interstitial transport. We demonstrate the broad utility of this approach by using type I collagen as the base hydrogel scaffold and probed for the biophysical effects of matrix cross-linking by human recombinant tissue transglutaminase II (hrTGII) and addition of exogenous HA. Surprisingly, the addition of hrTGII was seen to modify matrix structure and transport but not mechanical parameters. Furthermore, changes in matrix structure due to hrTGII were seen to be dependent on the concentration of collagen. Supplementation of HA at different collagen concentrations altered matrix microstructure and mechanical indentation but not transport parameters. Collectively the results from this study demonstrate intriguing effects for these modifications made possible through the advantages enabled by the reported characterization approach.

## 2. Materials and Methods

### 2.1 Preparation and casting of collagen matrices

Polymerized collagen matrices were prepared per manufacturer’s instructions. Briefly, rat tail type I collagen stored in acidic solution (Corning Life Sciences) was neutralized to pH = 7.4 using sodium hydroxide in 10X phosphate buffered saline. DMEM media was then utilized to adjust the final concentration of collagen to 3 and 6 mg/ml. Collagen gels were pre-incubated at 4°C for approximately 12 min prior to casting to enhance fiber formation^24^. Human recombinant tissue transglutaminase II (hrTGII) (R&D Systems) was added to collagen matrices by dissolving this component (1.0 μg) into DMEM during hydrogel preparation^26^. hrTGII drives a transamination reaction that produces amide cross-links between glutamine and lysine residues in collagen fibrils^27^. Collagen cross-linking by hrTGII was verified by primary amine group quantification with trinitrobenzene sulfonic acid^27^, as previously described. Hyaluronic acid (HA) salt from bovine vitreous humor (Sigma, MW ~400 kDa^28^) was dissolved in DMEM and added during the casting of collagen matrices^29^. Incorporation of HA was verified by alcian blue staining^30^ **(Supplementary Figure 1)**.

Upon neutralization and pre-incubation, collagen gels were pipetted into the different configurations used for the characterization scheme **(Fig. 1)**. For indentation testing, 300 μL of gel was casted in a cylindrical mold obtained from laser cutting a 48 well plate (Eppendorf). For mass transport measurements (i.e. hydraulic permeability and diffusivity) and confocal reflectance imaging of collagen fibers, gels were casted inside polydimethylsiloxane (PDMS) microchannels bonded irreversibly to a glass coverslip. Upon casting, gels were placed in a humidified incubator at 37°C for at least 20 min to complete polymerization before being subsequently hydrated with DMEM media. All samples were stored at 37°C prior to data acquisition.

**Figure 1.**
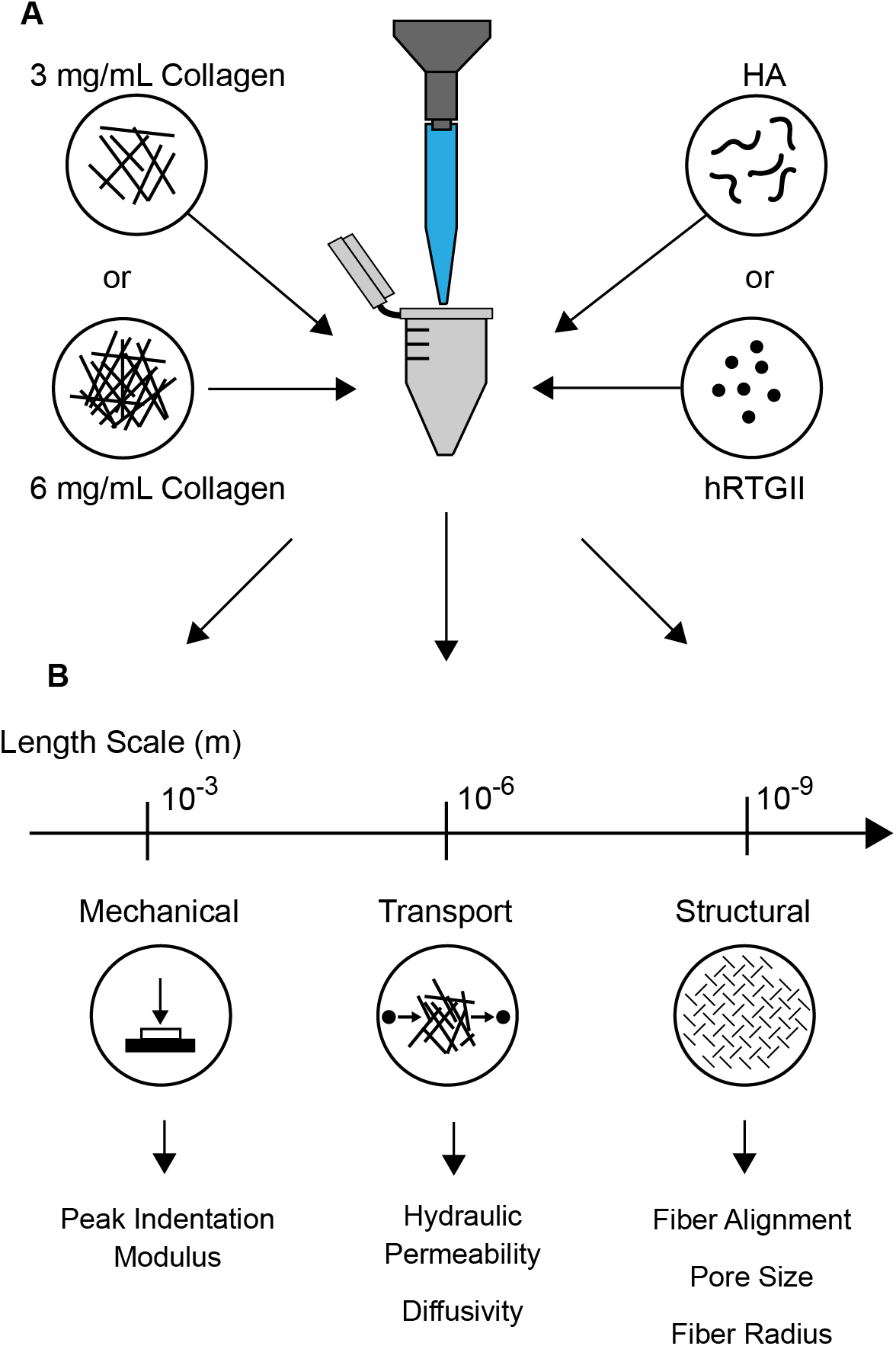
Integrated approach for characterizing collagen-based matrices supplemented with either hrTGII or HA. **A)** Collagen gels were prepared and characterized **(B)** with measurements of mechanical, transport, and structural parameters across varying length scales.

### 2.2 Mechanical indentation testing and data analysis

The mechanical stiffness of different ECM hydrogels was represented by the peak indentation modulus, which is a measure of the resistance to peak compressive loading^31^. Indentation testing of collagen-based matrices was conducted as previously described^31^ **(Fig 2A-C)**. Briefly, the well containing the gel was loaded into a high precision mechanical load frame (ElectroForce 5500, TA Instruments) and subjected to four incremental stress-relaxation indentations at a strain rate 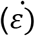 of 10 percent of the gel thickness (t_gel_) using a metallic circular flat ended indenter **(Fig. 2A-B)**. Each indentation step was performed to a depth of ten percent of the sample thickness (~3 mm). Peak indentation loads were then converted to peak indentation stresses using known indenter geometry and plotted against the corresponding strain. The peak indentation modulus was then estimated by computing the slope of the peak indentation stress-strain curve produced from these measurements **(Fig. 2C)**. Post-measurement analyses were done automatically using custom scripts written in MATLAB.

**Figure 2.**
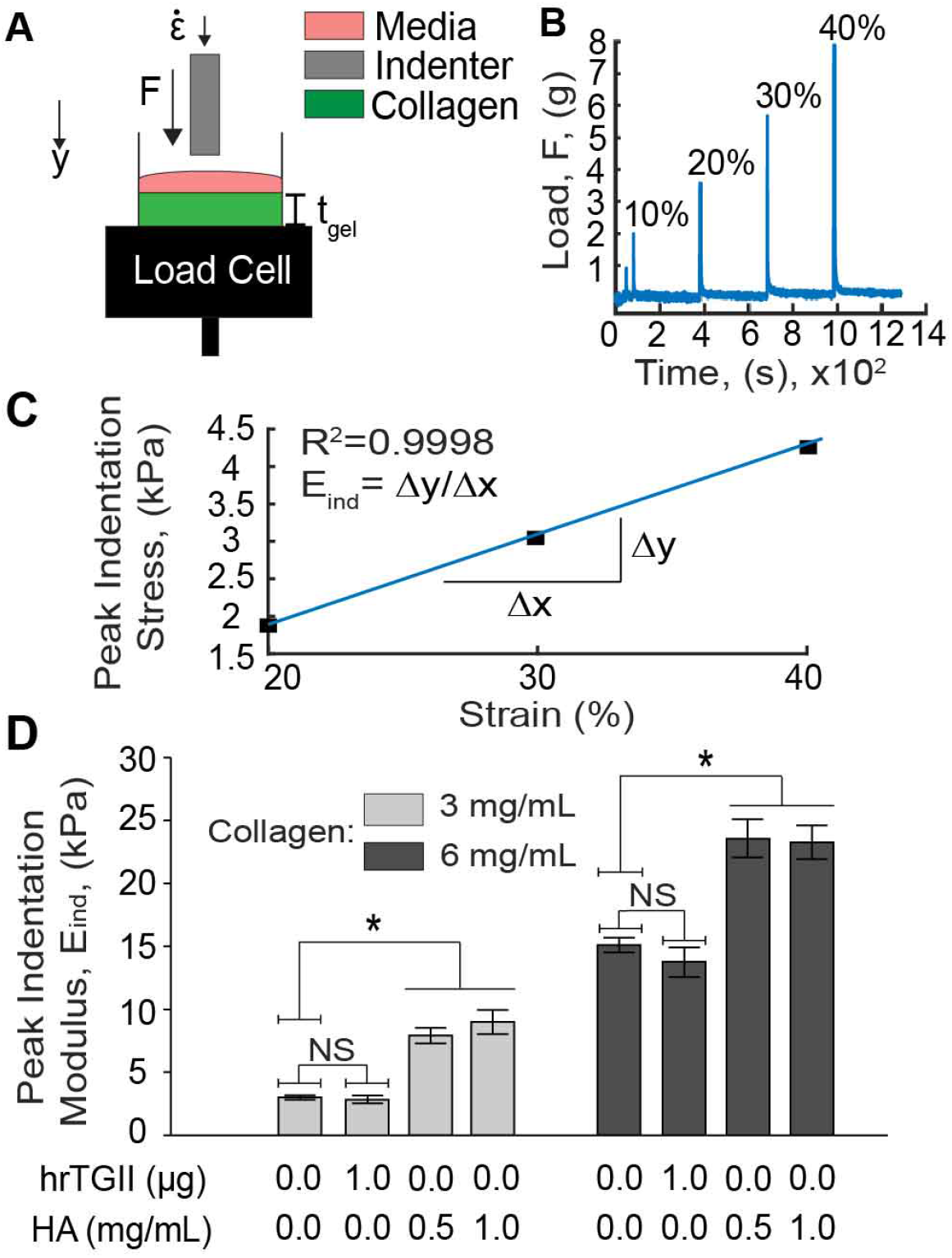
Quantification of peak indentation modulus of different ECM compositions. **A)** Collagen gels in cylindrical molds were loaded into a mechanical load frame and subjected to incremental indentation-relaxation steps at a strain rate 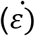 of ten percent of the gel thickness (*t*_*gel*_). **B)** The load applied to the indenter was recorded and plotted over time to obtain the peak indentation loads. **C)** The peak indentation loads were converted to peak indentation stresses and plotted against the corresponding indentation strain (starting from 20%) to compute the peak indentation modulus (*E*_*ind*_). **D)** Peak indentation modulus measurements. hrTGII did not alter the peak indentation modulus compared to control conditions. Addition of HA significantly increased the peak indentation loading for both 3 and 6 mg/ml collagen gels.

### 2.3 Microfluidic hydraulic permeability measurement

Convective transport of the collagen gels was characterized by the parameter hydraulic or Darcy permeability, which determines the flow rate in a porous material under an applied pressure gradient^32^. Hydraulic permeability was measured in a rectangular microfluidic device (Length: 5 mm; Width: 500 μm; Height: 1 mm) 48 hours post seeding as described in Hammer et al.^33^ Briefly, a height difference across the collagen gel was created by inserting a trimmed pipet tip at one of the ports and filling the tip with cell culture medium. This height difference produces a pressure gradient across the microchannel that generates flow through the seeded collagen type I matrix. The average velocity of the flow is estimated by tracking the motion of a fluorescent tracer dye within the microchannel (TRITC-BSA, MW = 65.5 kDa) using time lapse microscopy. Images were recorded every 5 seconds for a duration of 20-30 min using a Nikon TS-100F microscope equipped with a Q-Imaging QIClick camera controlled with NIS-Elements software. A custom MATLAB algorithm was then developed to automatically estimate the hydraulic permeability from the time lapse movies. The algorithm imports each frame and estimates the position of the dye profile by subtracting the image at t = i with the image at t = i−1 and tracking the centroid of the resulting difference image **(Supplementary Image 2)**. The position of the centroid is then plotted over time and the slope is obtained to approximate the bulk dye velocity **(Fig. 3)**. Darcy’s law was then used to calculate the hydraulic permeability.

**Figure 3.**
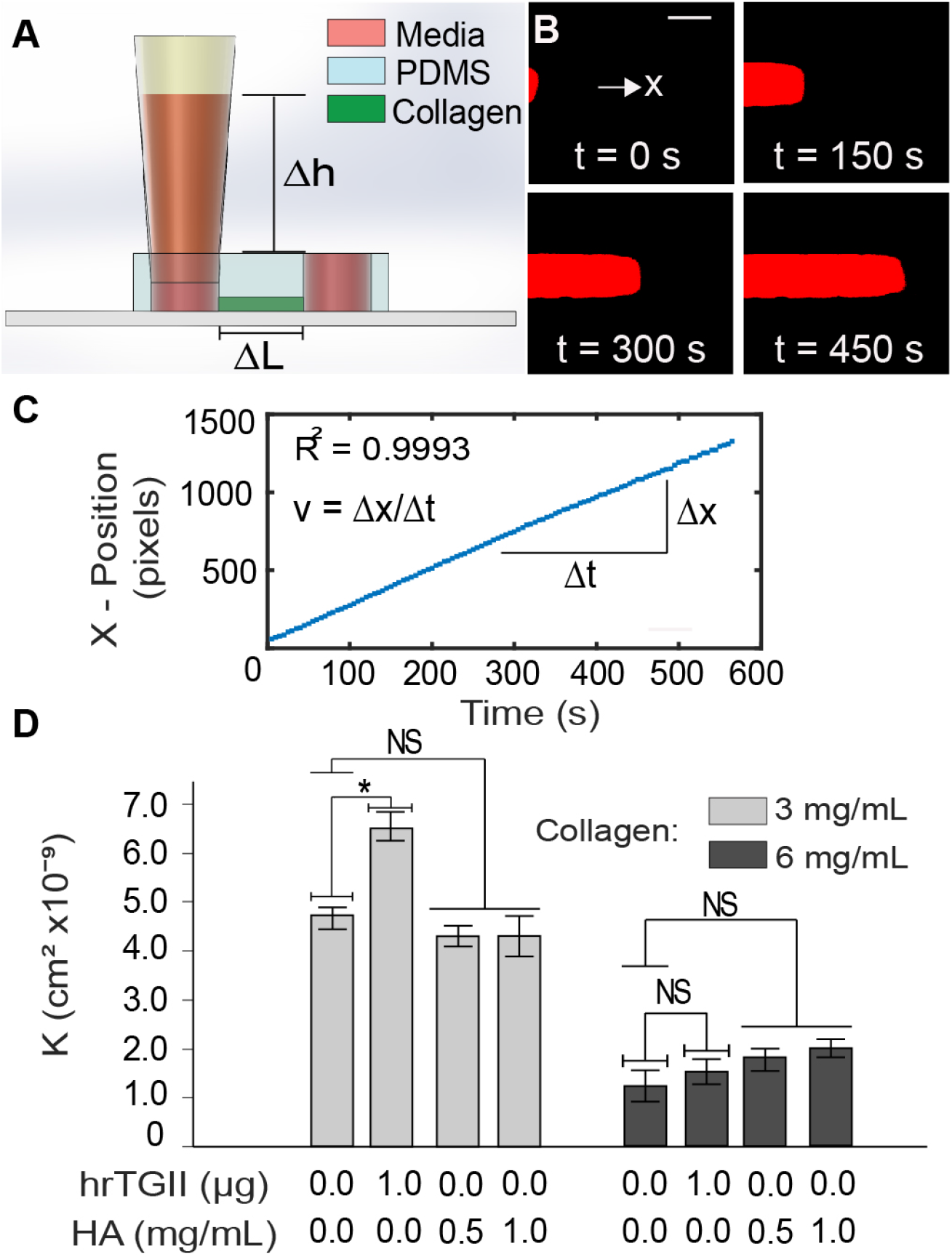
Hydraulic permeability measurements. **A)** Straight channel PDMS devices with 4 mm ports are sealed into a glass slide or coverslip and a trimmed pipet tip inserted (inset) to create a pressure difference that generates flow across the channel. Collagen type I of different compositions are injected and polymerized within the microfluidic device. **B)** Time lapse microscopy is used to track BSA-TRITC dye across the channel to obtain the average velocity of the flow **(C)** (Scale Bar = 150 μm). Darcy’s law is then used to estimate the hydraulic permeability. **D)** Hydraulic permeability quantification. hrTGII did increased the hydraulic permeability for 3 mg/ml collagen matrices but not for 6 mg/ml. Addition of HA did not alter the hydraulic permeability of the matrices for both collagen concentrations.

### 2.4 Measurement of diffusivity in microfluidic device

Quantification of matrix diffusivity to the fluorescent tracer dye (TRITC-BSA, MW = 65.5 kDa) was done with the use of Fluorescence Recovery After Photobleaching (FRAP) on rectangular microchannels (L: 20 mm, W: 3 mm, T: 20 μm)^34^. FRAP measurements were conducted on an Olympus FV1000 Multiphoton microscope equipped with a DeepSee MaiTai titanium-sapphire laser using a 40× water immersed objective (NA: 0.80) and an excitation wavelength of 840 nm. The laser was focused on a 40 μm circular region of interest (ROI) for 10 seconds to bleach the dye followed by image acquisition at intervals of 0.243 seconds for a total of 14 seconds **(Fig. 4A)**. Diffusivities were estimated using an automated custom made MATLAB script with a procedure previously described by Brancato et al.^35^ Briefly, the mean intensity inside the ROI was measured and converted to normalized fractional intensity (f). This parameter was then plotted versus time and fitted with an exponential curve to determine the half-recovery time τ where f = 0.5 (**Fig. 4B**). The diffusivity was then calculated by (1):

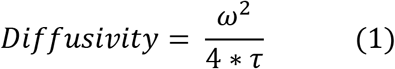

where *ω*^2^ is the radius of the bleach region (40 μm).

**Figure 4.**
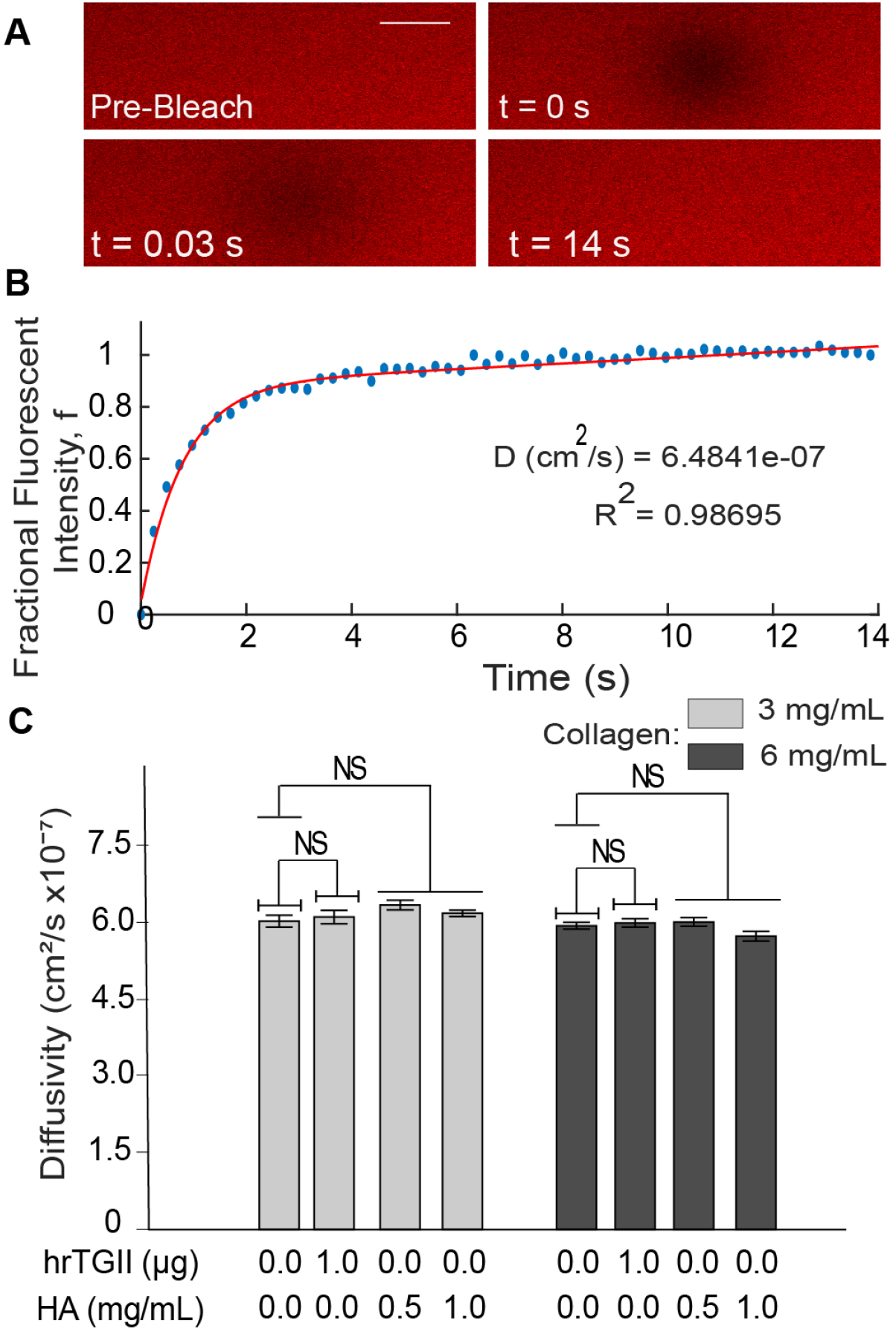
Interstitial diffusivity measurements with fluorescence recovery after photobleaching (FRAP). **A)** Images were recorded in the collagen-based matrices in PDMS microchannels loaded with BSA-TRITC dye. A bleached circular region was created containing and allowed to recover over time. (Scale Bar = 40 μm) **B)** The intensity in the bleached region is quantified over time and used to calculate the fractional fluorescent intensity. This parameter is plotted over time and fitted to estimate the diffusivity of the matrix. **C)** Diffusivity quantification. Modifications to collagen matrices with hrTGII or HA did not significantly alter the diffusivity for both collagen concentrations considered.

### 2.5 Quantification of Peclet number

The relative contributions of diffusion and convection to mass transport in matrices were quantified via the dimensionless parameter, Peclet number (Pe)^36^:

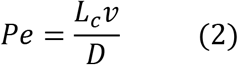

Where *L*_*c*_ denotes the characteristic length of interstitial transport (~100 μm), *v* denotes the fluid velocity through the matrices measured in Section 2.3 for estimating hydraulic permeability, and *D* denotes the matrix interstitial diffusivity.

### 2.6 Confocal reflectance microscopy image acquisition

Collagen fibers within the microfluidic device (**Fig. 5A**) were imaged using confocal microscopy on a Nikon A1R Live Cell Imaging Confocal Microscope via a 40× 1.3 NA oil immersion lens controlled with NIS-Elements Software. The reflectance signal was detected by exciting samples with a 487 nm laser, passing the reflected light through a 40/60 beam splitter, and collecting it in a detector for visualization of collagen fibers^37^. Confocal reflectance stacks of approximately 50 μm in height with a z-step of 0.59 μm were acquired for image analysis (80-90 total images per stack). In each experiment, duplicate devices were imaged for each condition. When imaging, 3 stacks per devices were obtained. Each condition was run 3 to 5 times in independent experiments.

**Figure 5.**
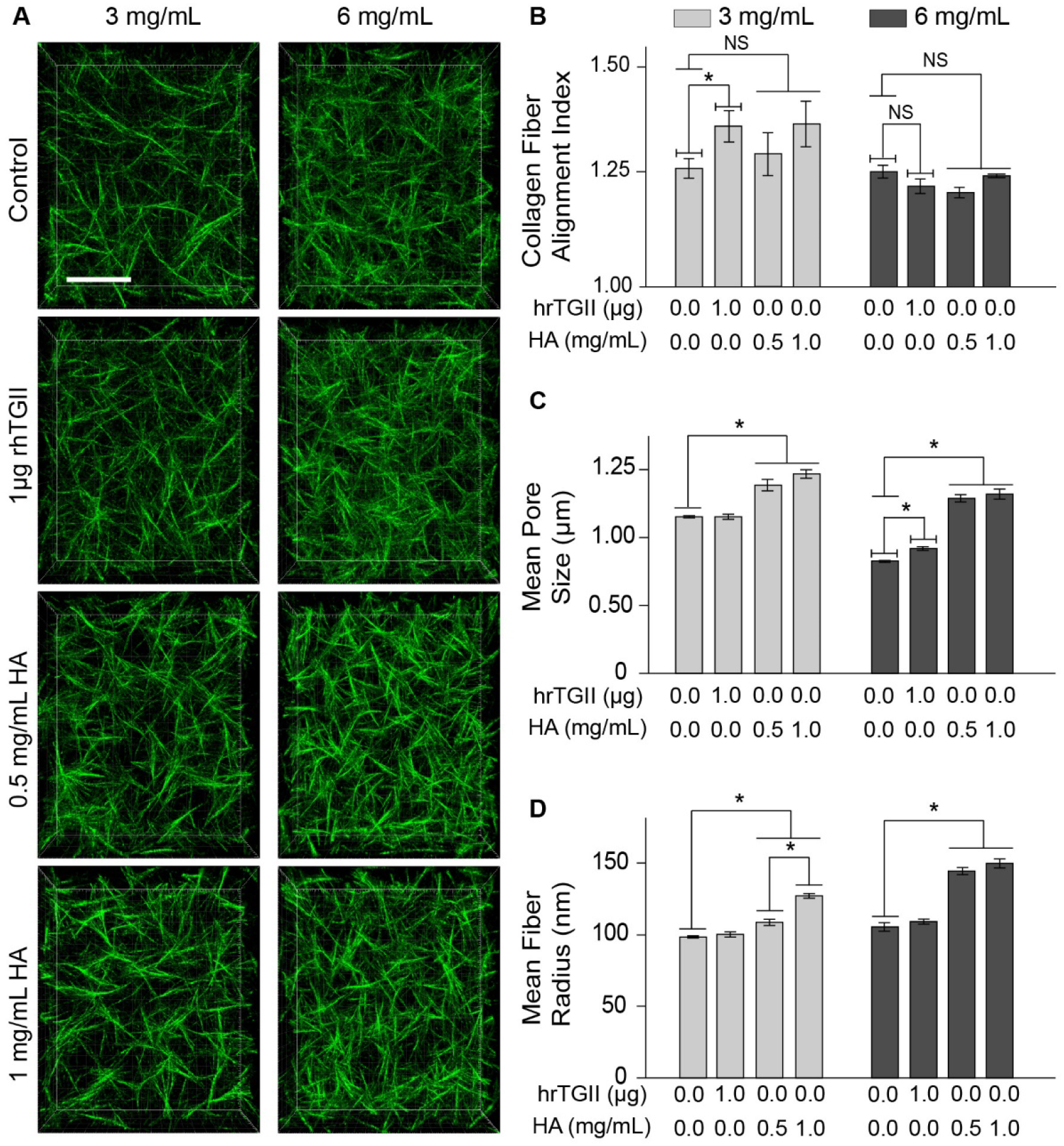
Microstructural characterization of collagen-based matrices. **A)** Confocal projections of 3 and 6 mg/ml collagen matrices with added hrTGII and HA (Scale Bar = 20 μm). **B)** Average matrix pore size determined by the nearest obstacle method. hrTGII did not alter the pore size of 3 mg/ml collagen matrices and slightly increased the pore size of 6 mg/ml collagen matrices. HA increased the average matrix pore size for both collagen concentrations. **C)** Alignment of collagen fibers quantified with alignment index. hrTGII increased the alignment of 3 mg/ml collagen matrices but not for 6 mg/ml matrices. Addition of HA did not alter the alignment index at either collagen concentration. **D)** Mean fiber radius quantified through 3D skeletonization. Addition of HA significantly increased the radius of collagen fibers at both collagen concentrations. hrTGII did not alter the radius of the collagen fibers.

### 2.7 Collagen fiber image analysis

Confocal reflectance stacks of collagen fibers were analyzed to quantify the fiber alignment, average pore size of the matrix, and fiber radii. Collagen fiber alignment was assessed using Fast Fourier transform (FFT) analysis as described previously^38^. Confocal reflection images of collagen fibers were converted into grayscale and analyzed for alignment using a custom-made MATLAB program. Alignment was summarized with the computation of an alignment index that indicates the degree of anisotropy of the collagen fibers by comparing intensity fractions within 20° of the most occurrent angle in the image to the same fraction in a random histogram. With this criterion, values close to 1 represent randomly aligned fibers while values greater than 1 represent highly aligned fibers. The average pore size of the matrix was estimated by use of the Nearest Obstacle Distance (NOD) Method^39^. In this approach the average pore size was quantified by fitting a Rayleigh distribution to the Euclidian distance transform of the binarized stack to obtain the biased average pore radius of the network. This value is then corrected for missing out of plane fibers, assuming a cut off angle of 43 degrees. To quantify fiber radii, stacks were imported into FIJI and subsequently skeletonized in 3D using the Skeleton tool. The total skeleton length was then computed and used to estimate the radius of the fibers from the known collagen concentration^40^.

### 2.8 Statistical analysis

Differences in properties of hrTGII supplemented collagen hydrogels were compared to control matrices at each concentration with the use of a t-test. To evaluate the differences between control matrices and those with added HA, ANOVA was used followed by Tukey’s post testing for pairwise comparisons between collagen controls and different concentrations of HA. All data is presented with bar graphs alongside error bars representing the standard error from the mean. Each condition was characterized independently at least three times and all data pooled together for statistical evaluation. A p value of less than 0.05 was used as a threshold for statistical significance for the differences observed between conditions.

## 3. Results

### 3.1 Differential effects of TGII and HA on altering the mechanical stiffness of collagen-based matrices

We first determined the bulk mechanical properties of different ECM compositions by measuring the peak indentation modulus **(Fig. 2)**. This parameter is dependent on both fibrillar and non-fibrillar matrix components, thus providing a rationale to assess how hrTGII-mediated cross-linking of fibrils and HA modify the mechanical behavior of collagen-based matrices. In addition, hrTGII and HA have been used to manipulate cell behavior in hydrogels, but their impact on altering matrix mechanical stiffness have not been reported. For the collagen-only matrix, at 3 mg/ml, the indentation stiffness was 3.01 kPa, and at 6 mg/ml, the indentation stiffness was 14.8 kPa **(Fig. 2D)**. For some physiological context, the stiffness profile for normal breast tissue is ~3.00 kPa^14, 41^ while the previously reported range of stiffness measurements for breast cancer tissue is 10.0-42.0 kPa^42^. Our results demonstrate that 3 mg/ml collagen recapitulates the tissue stiffness profile for normal breast while 6 mg/ml approximates the lower range of stiffness for breast cancer.

Next, we measured the indentation modulus of collagen that was cross-linked with hrTGII. At 3 mg/ml collagen, addition of hrTGII resulted in an indentation modulus of 2.87 kPa; at 6 mg/ml collagen, addition of hrTGII resulted in an indentation modulus of 14.3 kPa. Importantly, collagen cross-linking by hrTGII was verified by primary amine group quantification with trinitrobenzene sulfonic acid^27^ **(Supplementary Figure 1)**. Therefore, cross-linking by hrTGII application did not significantly modify the indentation stiffness of collagen gels for both concentrations considered **(Fig. 2D)**. In contrast to hrTGII, addition of HA to the collagen gels significantly increased the peak indentation modulus for both collagen concentrations considered **(Fig. 2D)**. At 3 mg/ml of collagen, addition of HA at concentrations of 0.5 mg/ml and 1.0 mg/ml significantly increased the indentation stiffness compared to collagen-only matrices to 7.92 kPa and 9.00 kPa respectively. At 6 mg/ml of collagen, addition of HA (0.5 mg/ml and 1 mg/ml) significantly increased the indentation stiffness to ~23 kPa for both HA concentrations. Given the ability of HA to withstand compressive loading in tissue such as cartilage^31^, this increase in indentation stiffness is expected, and can be attributed to HA affecting the ability of collagen fibers to reorganize in response to the applied indentation load^43^. We verified that increased HA concentration in the collagen gels resulted in increased alcian blue staining **(Supplementary Figure 1)**. Interestingly, increasing the concentration of HA from 0.5 to 1 mg/ml did not further enhance the indentation modulus, suggesting that this effect saturates beyond a certain HA concentration in collagen-based matrices.

### 3.2 Effects of TGII and HA on the convective and diffusive transport properties of collagen-based matrices

Next, we characterized the effects of hrTGII and HA on mass transport by convection and diffusion in collagen-based matrices **(Supplementary Table 1)**. For these studies, we introduced the same ECM compositions used for the indentation testing into a single channel microfluidic device. To characterize convection through the 3-D ECM, we measured the hydraulic or Darcy permeability to fluid flow (see Material and Methods). Hydraulic permeability is known to be dependent on ECM structure, composition, and concentration^44^. Therefore, as expected, increasing the collagen concentration from 3 mg/ml to 6 mg/ml significantly decreased the hydraulic permeability by 74% (from 4.72×10^−9^ cm^2^ for 3 mg/ml to 1.22×10^−9^ cm^2^ for 6 mg/ml) **(Fig. 3D)**. However, addition of hrTGII resulted in a significant increase of 38% in hydraulic permeability (from 4.72×10^−9^ cm^2^ to 6.51×10^−9^ cm^2^) compared to control at 3 mg/ml collagen but not at 6 mg/ml concentration **(Fig. 3D)**. Interestingly, supplementation of HA to the collagen matrices did not significantly alter the hydraulic permeability for both collagen concentrations considered **(Fig. 3D)**. At 3 mg/ml of collagen, HA addition resulted in hydraulic permeability values of ~4.30×10^−9^ cm^2^ for both HA concentrations. At 6 mg/ml of collagen, addition of HA at 0.5 mg/ml and 1 mg/ml resulted in slightly higher permeability values compared to control (1.81×10^−9^ cm^2^ and 2.00×10^−9^ cm^2^) but was not statistically significant.

We also measured the interstitial diffusivity, which like hydraulic permeability, is dependent on ECM properties but can also depend on the size and geometry of the tracer molecule^45^. In contrast to the hydraulic permeability measurements, we observed no significant difference in the average diffusivity for all of the ECM compositions tested using FRAP-based measurements **(Fig. 4C)**. In addition, the measured values for diffusivity (~6×10^−7^ cm^2^/s) in our *in vitro* setting were comparable to the ones previously recorded FRAP measurements *in vivo* (1×10^−7^ — 6×10^−7^ cm^2^/s). Simultaneous quantification of transport parameters within 3-D matrices enabled an estimation of the Peclet number (Eq. 2), which is the dimensionless ratio of convection relative to diffusion **(Supplemental Table 1)**. A Peclet number greater than 1 indicates convection-dominated transport while a number less than 1 indicates diffusion-dominated transport. For macromolecules comparable to BSA and nanoparticles, convection is normally the dominant transport mechanism in tissue *in vivo*^46^. Taking the length scale in Eq. 2 to be 100 μm (or the average interstitial distance between blood vessels in normal tissue^46^) the Peclet numbers were approximately 30 and 10 for 3 and 6 mg/ml collagen. Addition of hrTGII to collagen at 3 mg/ml concentration increased the Peclet number by 40% (from 30 to 42). In contrast, addition of HA had no effect on the Peclet number. Collectively, these results demonstrate that convection is the dominant transport mechanism for BSA along a typical interstitial path in the reconstituted collagen-based matrices tested.

### 3.3 Alternations to the collagen matrix microstructure by TGII and HA

Next, we assessed modifications to the collagen matrix microstructure with confocal reflectance microscopy **(Fig. 5A)**. These images were then analyzed with different image processing schemes to compute three parameters for the collagen network: 1) alignment index (dimensionless), 2) mean matrix pore size (μm), and 3) mean fiber radius (nm). In addition, the modifications to these microstructural parameters can be correlated with the bulk mechanical **(Fig. 2)** and transport properties **(Figs. 3 and 4)** of the different ECM compositions. The alignment index, mean pore size, and mean fiber radius measurements for 3 mg/ml were 1.15, 1.15 μm, and 98.5 nm, respectively. For 6 mg/ml, the value for these parameters were 1.23, 0.83 μm, and 105 nm. As expected, increasing the collagen concentration from 3 mg/ml to 6 mg/ml decreased the mean pore size but did not affect either alignment or mean fiber radius^40, 47^.

We then compared the effects of hrTGII and HA with collagen-only controls on matrix microarchitecture parameters. Addition of hrTGII increased the alignment of collagen fibers at 3 mg/ml by 8% (from 1.27 to 1.36) but not at 6 mg/ml collagen concentration **(Fig. 5B)**. In contrast, supplementing the collagen matrices with HA did not significantly alter the alignment of fibers for both collagen concentrations considered. For the mean matrix pore size measurements, addition of hrTGII had no effect at 3 mg/ml collagen but exhibited modest increases at 6 mg/ml (from 0.83 μm to 0.92 μm) with statistical significance **(Fig. 5C)**. However, it must be noted that the estimated pore size for the collagen-only control and hrTGII supplemented matrices at 6 mg/ml collagen lies below the resolution limit for the NOD (~1 um)^39–40^. Therefore, the actual pore sizes for the 6 mg/ml collagen matrices may be smaller and the difference in pore sizes between the two collagen concentrations evaluated may be greater. In contrast to the hrTGII supplemented matrices, addition of HA significantly increased the mean pore size for both 3 mg/ml (by 30%, from 1.15 um to 1.39 um and 1.47 um) and 6 mg/ml (by 57%, from 0.83 um to ~ 1.30 um for both HA conditions) collagen concentrations. The increase in matrix pore size may be attributed to HA-induced swelling, an effect that has been previously observed when incorporating HA in collagen gels^48–49^. Finally, for mean fiber radius, addition of hrTGII had no effect. Conversely, addition of HA significantly increased the mean fiber radius for both 3 mg/ml (by 10-30%, from 98.5 nm to 109 nm (0.5 mg/mL HA) and 127 nm (1 mg/mL HA) and 6 mg/ml (by 43%, from 105 nm to 148 nm for both HA conditions) collagen concentrations. **(Fig. 5D)**. This outcome may be explained by previous reports of HA associating around the collagen fibers during collagen fibrillogenesis^50^.

## 4. Discussion

Understanding the relationship between ECM composition and biophysical properties is necessary to help advance the utility of *in vitro* tissue-equivalent systems for studying cell behavior. This study puts forth a versatile, robust, and accessible experimental characterization scheme for interstitium mimicking materials based on indentation testing, quantification of transport via microfluidics, and confocal reflectance microscopy analysis. Here, we carefully characterized how hrTGII mediated cross-linking and HA regulate the physical characteristics of reconstituted acellular collagen-based matrices. TGII was selected as a representative example of enzymatic cross-linking that is upregulated in pathologies such as idiopathic pulmonary fibrosis^3^. HA was chosen as a representative example due its prominent role in fibrotic and desmoplastic diseases^51^.

Collagen cross-linking has been correlated with increases in stiffness in the context of cell seeded matrices. For instance, it has been reported that hrTGII may indirectly modulate the stiffness of cell-seeded collagen matrices by altering cellular responses^52^ such as contractility^53^ that are independent of its cross-linking functions. Yet, our measurements in acellular collagen-based matrices demonstrate no modification of indentation stiffness due to cross-linking by hrTGII alone. Therefore, our results support previous findings that direct changes to the mechanical integrity of fibrous collagen scaffolds require the formation of additional cross-link products than the ones enabled by hrTGII application under the provided conditions^54^. With regards to matrix transport properties, solute diffusivity was unaffected by any of the modifications considered, including hrTGII application. In contrast, hrTGII significantly increased the hydraulic permeability for 3 mg/ml collagen gels whereas it did not modify hydraulic permeability of 6 mg/ml collagen gels. We also observed significantly increased fiber alignment due to hrTGII treatment for 3 mg/ml but not 6 mg/ml collagen gels. Thus, the alignment index computations of the collagen fibers help explain the outcomes for hydraulic permeability as fibers aligned parallel to the direction of flow confer less resistance than fibers aligned at an angle^55^.

In contrast to hrTGII, HA (MW ~400 kDa) supplementation noticeably increased the measured indentation stiffness of both 3 mg/ml and 6 mg/ml collagen gels. In addition, increases in the indentation stiffness for HA supplemented matrices were correlated with enhanced mean matrix fiber radius and mean matrix pore size. Both of these microstructural changes to the collagen network may be a function of the size or molecular weight of the HA molecules, which in turn can impact the chemical expansion or swelling properties^56^. While our study did not consider resistance to indentation due to the swelling stresses conferred by HA, this parameter can be studied using more specialized testing approaches^57^. Our results for mean fiber radius demonstrate that HA significantly increases the radii of collagen fibers for all the collagen/HA compositions evaluated. Moreover, these results also suggest that collagen fiber re-orientation to an applied peak indentation load may depend on the radius of collagen fibers.

With regards to transport properties, cell-mediated HA deposition had been correlated with decreased hydraulic permeability in cell-laden collagen hydrogels *in vitro* and *in vivo*^33, 55, 58–59^. However, in the context of this study, no modification to hydraulic permeability due to HA supplementation of acellular collagen matrices was observed. These results suggest that the flow reducing effects of HA on transport properties may be dependent on both its assembly and distribution in the collagen ECM^60^. For instance, it is believed that cells assemble high molecular weight HA (MW > 1000 kDa) and degrade it into fragments of varying lower molecular weights (20 monosaccharides to 1000 kDa), which can impart different biological functions^61^. Based on these criteria, we believe the HA used in this study (MW~400 kDa) is more representative of low molecular weight HA. Previous work on collagen and HA (MW~155 kDa) assembly *in vitro* suggest that HA associates around the collagen fibers as opposed to being distributed within the interfibrillar space^62^, thereby minimizing hindrance to fluid flow. Since our study was in acellular matrices *in vitro* and using a comparable molecular weight HA, we believe that our results point to the latter scenario. Moreover, our microstructural analysis by confocal reflectance microscopy showed that HA significantly increases both the mean fiber radii and matrix pore sizes for all of the collagen/HA combinations tested. When integrating our microstructural and transport results, it appears that low molecular weight HA-mediated increases in mean fiber radius and matrix pore size are in effect neutralizing each other in terms of transport path in porous media and subsequently hydraulic permeability. Furthermore, since HA had no effect on the hydraulic permeability of collagen matrices, this result suggests that the observed increases in indentation stiffness for this ECM composition were due to collagen fiber enhancement as opposed to increased pressurization of interstitial fluid. Finally, our results demonstrate that for both hrTGII and HA-modified matrices, mechanical and transport properties may act independent of each other and can be used as different measures to profile tissue or tissue-equivalent constructs.

In addition to hydraulic permeability, another important transport parameter is diffusivity. Our results were that diffusivity to BSA was unaffected by any of the ECM modifications considered, even when increasing the collagen content from 3 mg/ml to 6 mg/ml. This outcome suggests that at relative low collagen gel fractions, diffusive transport is unaffected when increasing the collagen content within the range of concentrations considered. However, the effect on diffusivity may be more pronounced when considering higher collagen concentrations than the ones used in our studies^56^. Our results also support the notion that hydrogel structure does not modify the diffusion of particles with a hydrodynamic radius (~3.5 nm, BSA) significantly smaller than the pore size of the matrix (1-1.25 μm)^63^. Given that the diffusivities for the tested matrices were comparable, changing the Peclet number of these matrices is achieved by modifying the hydraulic permeability.

For fibrillar scaffolds, establishing a relationship between transport efficiency with microstructural parameters has been challenging due to complex ECM network morphologies that can preclude straightforward measurements of transport parameters. However, this study demonstrates the utility of microfluidic devices for addressing this challenge as all the transport measurements and microstructural analysis were done in microfluidics devices integrated with collagen-based scaffolds. Pressure gradients in microfluidics devices can be specified with relative ease for accurate hydraulic permeability measurements^36^. Also, the optical clarity of the materials used for microfluidic device fabrication (i.e. PDMS channels sealed against a coverglass slide) are conducive for high-resolution confocal reflectance microscopy and FRAP-based diffusion measurements. We previously leveraged microfluidic systems to assess functional changes in hydraulic permeability mediated by cancer-associated stromal fibroblasts that were perturbed biologically by either genetic manipulations or targeted pharmacological inhibitors^33, 59^. Therefore, we foresee the integrated biophysical characterizations from this study to be part of the toolkit of functional assessments used in microfluidic-based organ-on-a-chip settings that can be readily scaled-up for high-throughput screening applications^64^.

While this study focused on characterizing acellular type I collagen, the physical profiling scheme presented in this study can also be applied to other natural and synthetic hydrogel materials used in tissue engineering applications (e.g. fibrin, matrigel, alginate) as well as assessing the effects of other matrix modifications during polymerization, such as temperature ramping. Moreover, future studies can be applied in cell-laden hydrogels to investigate the physical consequences due to various physiologically important cell-mediated matrix modifications, such as enzymatic degradation by matrix metalloproteinases, formation of more mature cross-links, and deposition of sulfated GAGs and other proteoglycans.

## 4 Conclusions

The characterization scheme presented in this study enabled new insight as to how hrTGII and HA modify the physical characteristics of reconstituted type I collagen matrices in the absence of cells. Given the widespread use of hydrogels across tissue engineering applications, another important aspect of this study is that it brings together a collection of methods that can assist others in decoupling matrix mechanics from matrix transport properties across different length scales. These advantages can be used by the biomaterials research community for biophysical profiling of interstitium mimicking materials. In addition, the results from this study can help guide future considerations of supplementing fibrous hydrogels with exogenous hrTGII and HA for tissue engineering applications by understanding the effects on structural, mechanical and transport parameters *in vitro*.

## Acknowledgements

JWS acknowledges NSF CAREER Award (CBET-1752106), The American Cancer Society (IRG-67-003-50), and The Ohio State University Materials Research Seed Grant Program, funded by the Center for Emergent Materials, an NSF-MRSEC, grant DMR-1420451, the Center for Exploration of Novel Complex Materials, and the Institute for Materials Research. AA, JCC, and JWS acknowledge funding from Pelotonia. MC-M acknowledges funding from Graduate Enrichment and Discovery Scholars Fellowships from Ohio State University and from a Graduate Diversity Supplement from the National Heart, Lung, and Blood Institute (R01HL141941-02S1). Confocal reflectance microscopy images presented in this report were generated using instruments and services at the Campus Microscopy and Imaging Facility, The Ohio State University. This facility is supported in part by grant P30 CA016058, National Cancer Institute, Bethesda, MD.

