## Supplementary Data for "Integrated biophysical characterization of fibrillar collagen-based hydrogels"

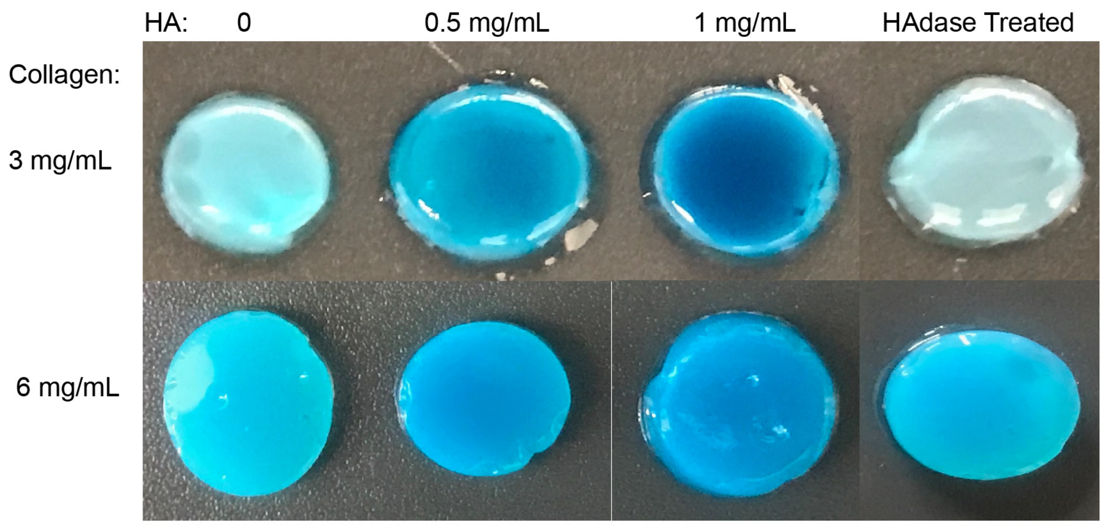


**B**

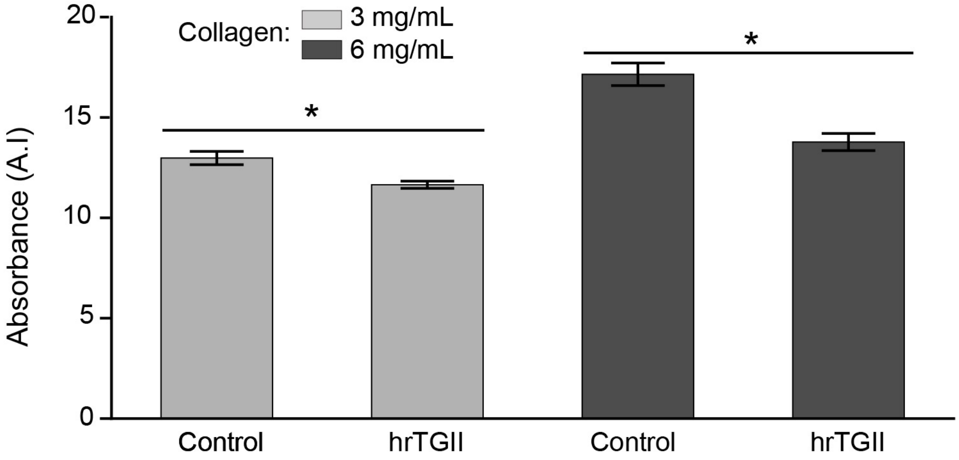


**Supplementary Figure 1. Modification of collagen matrices with HA and hrTGII. A)** Incorporation of HA into matrices was verified with increased alcian blue staining with increasing HA concentration. HA specificity was confirmed by pretreatment with hyaluronidase (HAdase) prior to straining. **B)** Collagen cross linking by hrTGII quantification. Cross linking was confirmed by reduction of free amine groups using absorbance of trinitrobenzene sulfonic acid at 420 nm (*p<0.05, t-test).

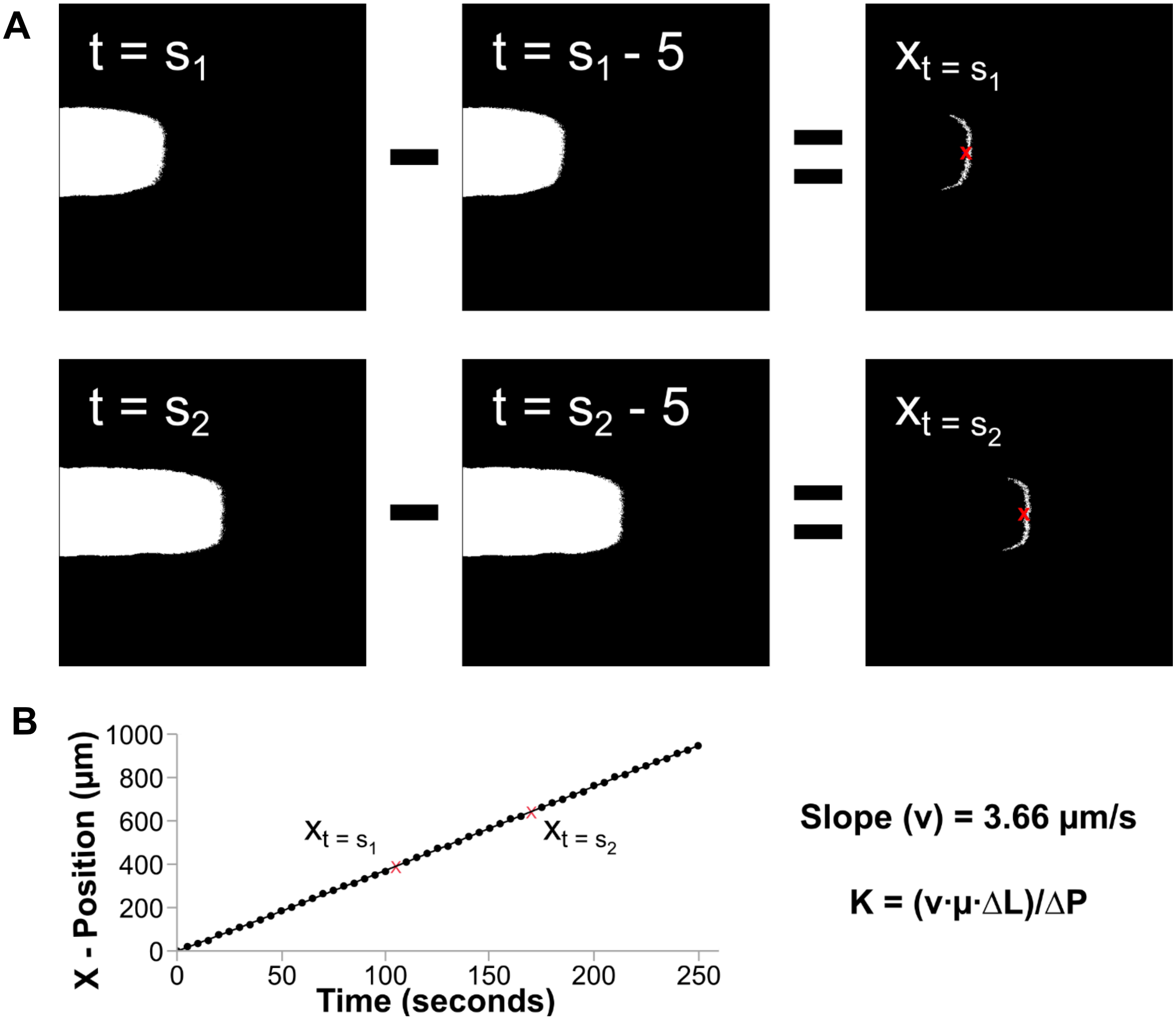


**Supplementary Figure 2. Tracking of dye flow profile for measurement of hydraulic permeability. A)** Time-lapse images are imported into MATLAB and binarized. The binarized images are then subtracted frame by frame and the resulting profile centroid x-position recorded. **B)** This position is plotted against time and the slope computed to determine the average flow velocity. Darcy's law is then used to calculate the hydraulic permeability.

| Collagen: | | $E_{indentation}$ (kPa) | K(cm² x$10^{-9}$) | $D_{BSA}$ (cm²/s x$10^{-7}$) | Pe |
|---|---|---|---|---|---|
| 3 mg/mL | Control | 3.01± 0.74 | 4.72± 0.28 | 6.32± 0.25 | 29 |
| | 1μg hrTGII | 2.87± 0.32 | 6.51± 0.31 | 6.17± 0.06 | 42 |
| | 0.5 mg/mL HA | 7.92± 0.61 | 4.30± 0.31 | 6.03± 0.12 | 28 |
| | 1.0 mg/mL HA | 9.00± 0.95 | 4.31± 0.28 | 6.32± 0.09 | 27 |
| 6 mg/mL | Control | 14.8± 0.61 | 1.22± 0.33 | 5.92± 0.07 | 8 |
| | 1μg hrTGII | 14.3± 1.19 | 1.51± 0.33 | 5.72± 0.10 | 10 |
| | 0.5 mg/mL HA | 24.7± 0.75 | 2.00± 0.28 | 5.98± 0.08 | 13 |
| | 1.0 mg/mL HA | 25.2± 0.51 | 1.81± 0.28 | 6.00± 0.08 | 12 |

**Supplementary Table 1. Summary of mechanical stiffness, hydraulic permeability, diffusivity, and peclet numbers for ECM compositions considered in this study.** Values denote the mean with the standard error of the mean use as bounds.
